# Neural signatures of the processing of temporal patterns in sound

**DOI:** 10.1101/261271

**Authors:** Björn Herrmann, Ingrid S. Johnsrude

## Abstract

The ability to detect regularities in sound (i.e., recurring structure) is critical for effective perception, enabling, for example, change detection and prediction. Two seemingly unconnected lines of research concern the neural operations involved in processing regularities: one investigates how neural activity synchronizes with temporal regularities (e.g., frequency modulation; FM) in sounds, whereas the other focuses on increases in sustained activity during stimulation with repeating tone-frequency patterns. In three electroencephalography studies with male and female human participants, we investigated whether neural synchronization and sustained neural activity are dissociable, or whether they are functionally interdependent. Experiment I demonstrated that neural activity synchronizes with temporal regularity (FM) in sounds, and that sustained activity increases concomitantly. In Experiment II, phase coherence of FM in sounds was parametrically varied. Although neural synchronization was more sensitive to changes in FM coherence, such changes led to a systematic modulation of both neural synchronization and sustained activity, with magnitude increasing as coherence increased. In Experiment III, participants either performed a duration categorization task on the sounds, or a visual object tracking task to distract attention. Neural synchronization was observed irrespective of task, whereas the sustained response was observed only when attention was on the auditory task, not under (visual) distraction. The results suggest that neural synchronization and sustained activity levels are functionally linked: both are sensitive to regularities in sounds. However, neural synchronization might reflect a more sensory-driven response to regularity, compared with sustained activity which may be influenced by attentional, contextual, or other experiential factors.

**Significance statement:** Optimal perception requires that the auditory system detects regularities in sounds. Synchronized neural activity and increases in sustained neural activity both appear to index the detection of a regularity, but the functional interrelation of these two neural signatures is unknown. In three electroencephalography experiments, we measured both signatures concomitantly while listeners were presented with sounds containing frequency modulations that differed in their regularity. We observed that both neural signatures are sensitive to temporal regularity in sounds, although they functionally decouple when a listener is distracted by a demanding visual task. Our data suggest that neural synchronization reflects a more automatic response to regularity, compared with sustained activity which may be influenced by attentional, contextual, or other experiential factors.

## Introduction

Natural sound environments are rich in statistical regularities and perceptual systems are sensitive to these regularities (Garrido et al., 2013; McDermott et al., 2013; Herrmann and Johnsrude, 2018). A regularity may be defined as reoccurring structure in sounds, and may include, among others, repeated patterns of brief tones or of a sound’s frequency modulation. Detection of regularities is crucial for efficient auditory scene analysis (Bendixen, 2014). It enables sensitivity to acoustic changes (Schröger, 2007; Winkler et al., 2009), allows generation of expectations about future sounds (Barnes and Jones, 2000; Jones et al., 2002; Herrmann et al., 2016), and supports perception of speech (Idemaru and Holt, 2011; Giraud and Poeppel, 2012; Peelle and Davis, 2013; Baese-Berk et al., 2014). In this paper, we investigate how regularity is represented in patterns of neural (electroencephalographic) activity.

Prior investigations of the neural mechanisms supporting the processing of regularities in sounds have largely been conducted along two seemingly unrelated lines (but see (Baldeweg, 2006; Winkler et al., 2009) for other measures). One line of research focuses on the propensity of neural oscillatory activity to synchronize with temporal regularities in sound environments (Lakatos et al., 2008; Nozaradan et al., 2011; Henry and Obleser, 2012; Lakatos et al., 2013b; Costa-Faidella et al., 2017; ten Oever et al., 2017). Temporally regular sound stimulation drives neural activity into the same rhythm as the stimulus rhythm. Synchronization of neural activity is thought to provide a listener with a reference frame that can be used to predict future sounds (Schroeder and Lakatos, 2009; Henry and Herrmann, 2014; Nobre and van Ede, 2018), which may, in turn, support perception of speech and music (Peelle and Davis, 2013; Doelling and Poeppel, 2015).

Another, more recent, line of research has revealed that the detection of repetition in a sequence of brief tones is marked by an increase in a sustained, low-frequency DC power offset in magneto-/electroencephalographic activity (Barascud et al., 2016; Sohoglu and Chait, 2016; Southwell et al., 2017). The sustained response increase appears independent of a specific type of structure: It has been observed for repeating sequences of tones with random frequencies (Barascud et al., 2016; Southwell et al., 2017), complex sounds made of isochronous tone sequences (Sohoglu and Chait, 2016), and coherent frequency changes in sounds made of broadband chords (Teki et al., 2016). Sustained response increases have been proposed to reflect a prediction-related increase in sensitivity when uncertainty is reduced (Barascud et al., 2016; Heilbron and Chait, in press).

The relation between neural synchronization and the sustained response for the detection of regularities is unknown. Neural synchronization to low frequency (~2–6 Hz) temporal regularities is strongest in auditory cortex (Herrmann et al., 2013; Keitel et al., 2017; Millman et al., 2017). Sustained response increases appear to be generated in a wide network of brain areas, including auditory cortex, hippocampus, parietal cortex, and inferior frontal gyrus (Barascud et al., 2016; Teki et al., 2016). Neural synchronization in sensory areas may provide the means to recognize the temporal regularity, which then is further processed in higher-order brain regions. However, it is unclear how neural synchronization and sustained response interact functionally.

In the current study, we conduct three electroencephalography (EEG) experiments to investigate the relation between neural synchronization and sustained neural activity. In Experiment I we test whether neural synchronization and sustained activity are simultaneously elicited by temporal regularity in sounds using a stimulation protocol similar to previous work (Barascud et al., 2016; Southwell et al., 2017). In Experiment II, we parametrically manipulate the degree of temporal regularity in sounds – and thereby the degree of neural synchronization – in order to investigate whether sustained activity is differently sensitive to temporal regularity compared to neural synchronization. Finally, using a selective attention paradigm, Experiment III tests the extent to which neural populations supporting neural synchronization versus sustained neural activity are modulated by attention, and thus the extent to which they may be potentially dissociable.

## General methods

### Participants

Participants were male and female students at the University of Western Ontario aged 17–29 years and with no reported neurological disease or hearing impairment. All participants had normal or corrected-to-normal vision and were naïve to the purposes of the experiment. Different participants were recruited for each of the three experiments. Participants gave written informed consent prior to the experiment, and either received course credit or were paid $5 CAD per half-hour for their participation. The experiments were conducted in accordance with the Declaration of Helsinki, the Canadian Tri-Council Policy Statement on Ethical Conduct for Research Involving Humans (TCPS2-2014), and were approved by the local Non-Medical Research Ethics Board of the University of Western Ontario (protocol ID: 106570).

### Procedure

All experimental procedures were carried out in a single-walled sound-attenuating booth (Eckel Industries). Sounds were presented via Sennheiser (HD 25-SP II) headphones and a Steinberg UR22 (Steinberg Media Technologies) external sound card. Stimulation was controlled by a PC (Windows 7, 64 bit) running Psychtoolbox in Matlab (R2015b).

Prior to the experiment, the hearing threshold for a 1600-Hz pure tone (Experiment I) or a narrowband noise centered on 1414 Hz (Experiments II and III) was determined for each individual using a method-of-limits procedure. The thresholds were used to present sounds during the EEG experiments at 55 dB above the individual sensation level. Thresholds were obtained by presenting sounds of 12-s duration that changed (either decreased or increased) continuously in intensity at 5.4 dB/s. Participants indicated via button press when they could no longer hear the tone (intensity decrease) or when they started to hear the tone (intensity increase). The mean sound intensity at the time of the button press was noted for 6 decreasing sounds and 6 increasing sounds (decreasing and increasing sounds alternated), and these were averaged to determine the individual hearing threshold.

### EEG recordings and preprocessing

EEG signals were recorded at 1024-Hz sampling rate from 16 (Ag/Ag–Cl-electrodes) electrodes and additionally from left and right mastoids (BioSemi, Amsterdam, The Netherlands; 208 Hz low-pass filter) while participants listened to sounds that either did or did not contain a regularity (for details on individual conditions see the experimental procedures below). Electrodes were referenced to a monopolar reference feedback loop connecting a driven passive sensor and a common-mode-sense (CMS) active sensor, both located posterior on the scalp.

Offline data analysis was carried out using MATLAB software (v7.14; MathWorks, Inc.). Line noise (60 Hz) was removed from the raw data using an elliptic filter. Data were re-referenced to the average mastoids, high-pass filtered at a cutoff of 0.7 Hz (2449 points, Hann window), and low-pass filtered at a cutoff of 22 Hz (211 points, Kaiser window). Data were down-sampled to 512 Hz and divided into epochs ranging from −1 to 5.8 s (Experiments I & II) or from −0.5 to 4 s (Experiment III). Epochs were shorter in Experiment III compared to Experiments I & II to accommodate differences in sound duration and trial structure (see below). Independent components analysis (runica method, (Makeig et al., 1996); logistic infomax algorithm, (Bell and Sejnowski, 1995); Fieldtrip implementation, (Oostenveld et al., 2011), RRID:SCR_004849) was used to identify and remove activity related to blinks and horizontal eye movements. Epochs that exceeded a signal change of 200 μV in any electrode were excluded from analyses. This pipeline was used to investigate neural synchronization (see below).

In order to investigate the sustained neural activity, the same pipeline was computed a second time, with the exception that high-pass filtering was omitted. Omission of the high-pass filter is necessary to investigate the sustained response, because the response is a very low-frequency signal reflecting a DC shift (Barascud et al., 2016; Southwell et al., 2017). Activity related to blinks and horizontal eye movements was removed using the identified components from the high-pass filtered data.

### EEG data analysis: Sustained response

Single-trial time courses were averaged for each condition (details on the conditions in each experiment are presented below). Note that previous studies that investigated the sustained response used a root-mean-square calculation instead of calculating the average across trials (Barascud et al., 2016; Sohoglu and Chait, 2016; Teki et al., 2016). The root-mean-square calculation results in only positive values, whereas the calculation of the average preserves information about polarity. Previous work utilized the root-mean square calculation because magnetoencephalography was recorded and absolute polarity in these recording cannot be interpreted. However, the polarity of the EEG signal may provide information about neural mechanisms as a negative-going wave may indicate an excitable state (Whittingstall and Logothetis, 2009). We therefore did not calculate the root-mean-square for the current EEG recordings but instead used the average across trials.

Separately for each electrode, response time courses were baseline corrected by subtracting the mean amplitude in the pre-stimulus time window (Experiments I & II: −1 to 0 seconds; Experiment III: −0.5 to 0 seconds; differences in the length are related to experiment-specific trial structure; see below) from the amplitude at each time point. In order to investigate the sustained response, signals were averaged across a fronto-central-parietal electrode cluster (Fz, F3, F4, Cz, C3, C4, Pz, P3, P4) and, for each condition, the mean amplitude was calculated within the time window during which a regularity could occur in sounds: 2–4.8 s (Experiment I), 2.2–4.8 s (Experiment II), 1.6–4 s (Experiment III). Selection of the fronto-central-parietal electrode cluster was motivated by the spatially wide-spread sustained response across electrodes (see Figures 1, 4, and 7) that is consistent with previous localizations of the sustained response in auditory cortex and higher-level brain regions (i.e., parietal cortex, hippocampus, inferior frontal gyrus) (Barascud et al., 2016; Teki et al., 2016).

**Figure 1:**
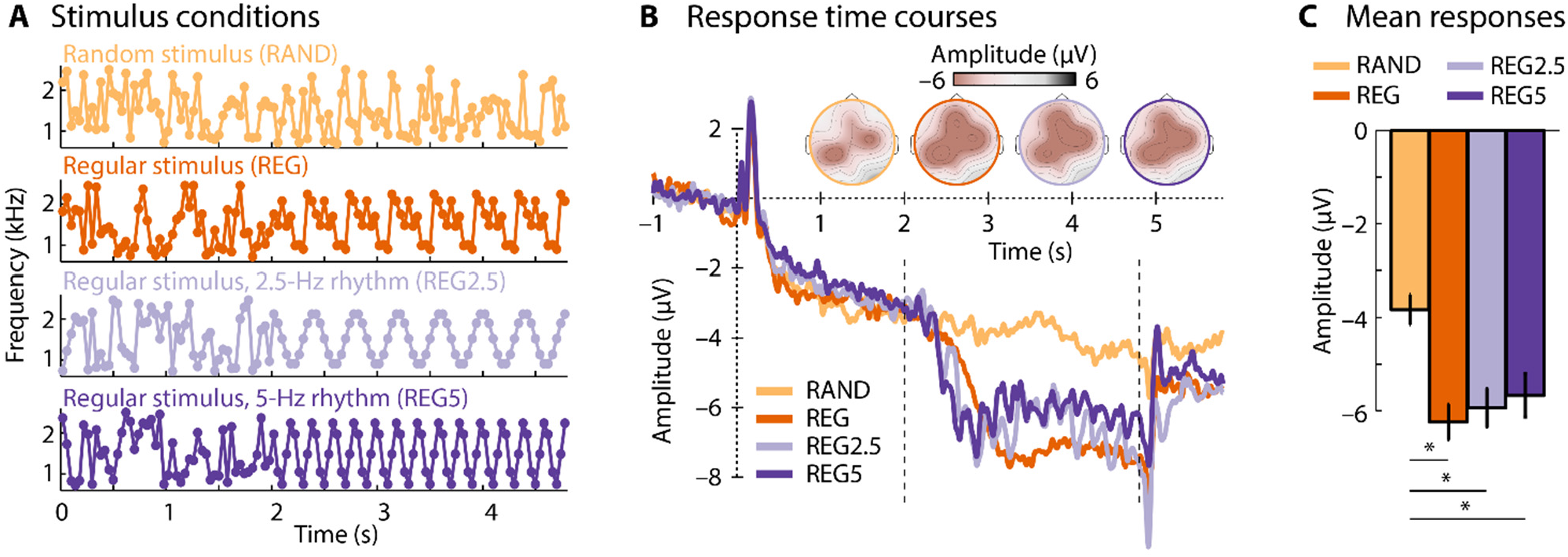
Stimulus conditions and neural responses for Experiment I. **A**: Frequency of brief tones (dots) making up a 4.8-s sound. An example for each stimulus condition is shown. **B**: Response time courses and scalp topographies for each condition. The dotted vertical lines mark the time window of interest used for analysis (2–4.8 s). **C**: Mean response magnitude for the 2–4.8 s time window. Error bars reflect the standard error of the mean (removal of between-subject variance; (Masson and Loftus, 2003). *p_FDR_ ≤ 0.05. Other stimulus effects (i.e., among the three REG sequences) were not significant.

**Figure 2:**
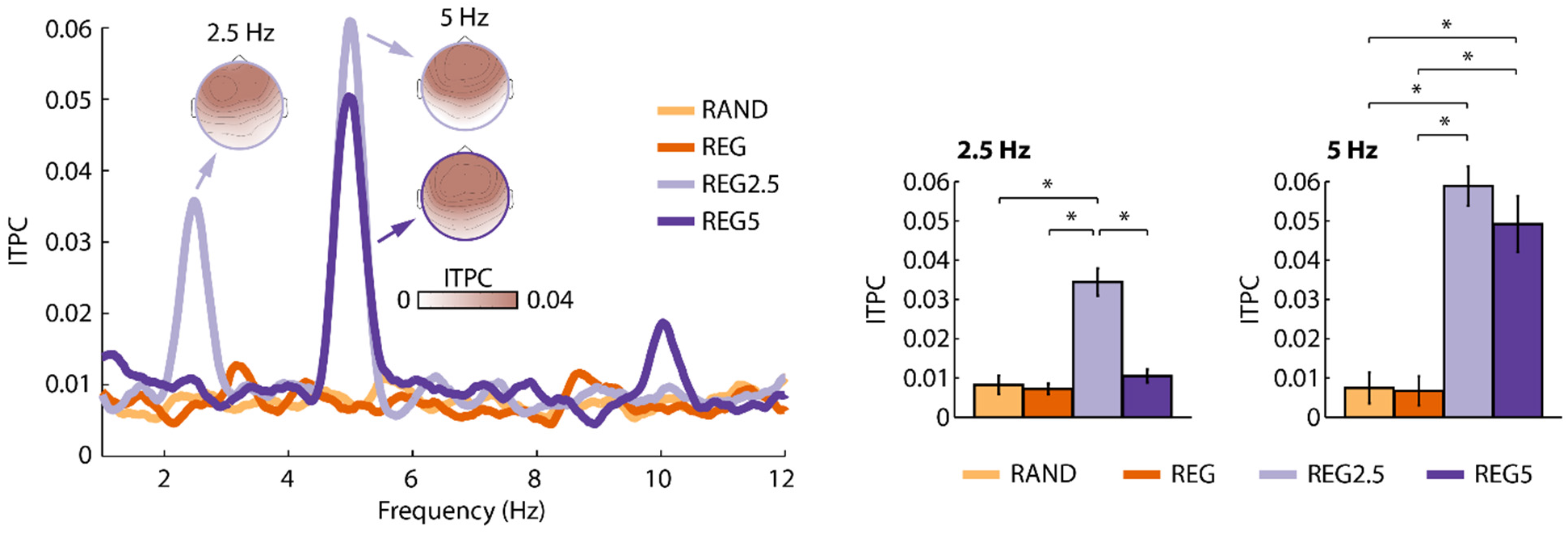
Results of the inter-trial phase coherence analysis for Experiment I. ITPC frequency spectrum (left) and ITPC values at the stimulus frequencies (right). Error bars reflect the standard error of the mean (removal of between-subject variance; (Masson and Loftus, 2003). *p < 0.05. Other stimulus effects were not significant.

### EEG data analysis: Neural synchronization

Neural synchronization was investigated using inter-trial phase coherence (ITPC) as a dependent measure (Lachaux et al., 1999). To this end, a fast Fourier transform (including a Hann window taper and zero-padding) was calculated for each trial and channel for the time window during which a regularity could occur in sounds: 2–4.8 s (Experiment I), 2.2–4.8 s (Experiment II), 1.6–4 s (Experiment III). The resulting complex numbers were normalized by dividing each complex number by its magnitude. ITPC was then calculated as the absolute value of the mean normalized complex number across trials. ITPC values can take on values between 0 (no coherence) and 1 (maximum coherence). ITPC was calculated for frequencies ranging from 1–12 Hz. For statistical analyses, ITPC was averaged across the fronto-central-parietal electrode cluster (Fz, F3, F4, Cz, C3, C4, Pz, P3, P4).

### Experimental design and statistical analysis

#### These experiments were not preregistered

Details of the critical variables and statistical tests for each experiment can be found below. In general, experimental manipulations were within-subject factors. Differences between experimental conditions were thus assessed using paired-samples t-tests, one-sample t-tests, or via repeated-measures analysis of variance (rmANOVA). Whenever the assumption of sphericity was violated, Greenhouse–Geisser correction was applied (Greenhouse and Geisser, 1959); we report the original degrees of freedom, the corrected p value, and the epsilon coefficient. False discovery rate (FDR) correction was applied to correct for multiple comparisons among post-hoc tests following an rmANOVA (Benjamini and Hochberg, 1995; Genovese et al., 2002). Correlations between conditions are reported as Spearman correlations and repeated-measures correlations (Bakdash and Marusich, 2017); using the ‘rmcorr’ implementation in Rstudio software; Version 1.1.419; https://www.rstudio.com/, RRID:SCR_000432). The repeated-measures correlation determines the common within-individual association for paired measures (two measures in Experiment I; four measures in Experiment II) for multiple individuals.

Throughout the manuscript, effect sizes are provided as partial eta-squared (η_p_^2^) when an analysis of variance (ANOVA) is reported and as r_e_ (r_equivalent_) when a t-test is reported (Rosenthal and Rubin, 2003). r_e_ is equivalent to a Pearson product-moment correlation for two continuous variables, to a point-biserial correlation for one continuous and one dichotomous variable, and to the square root of partial η^2^ for ANOVAs.

## Experiment I: Regularity detection – sustained activity and neural synchronization

Experiment I aimed to replicate previous observations of a sustained response elicited by a repeating sequence of brief tones (Barascud et al., 2016; Southwell et al., 2017), and, in addition, aimed to investigate whether neural synchronization co-occurs with changes in sustained activity levels.

### Participants

Sixteen participants took part in Experiment I (mean age [standard deviation]: 18.6 ±2 years; 8 female).

### Acoustic stimulation & procedure

Stimuli were similar to those used in previous studies (Barascud et al., 2016; Southwell et al., 2017). Stimuli were 4.8-s long sequences of 0.04-s tones (0.007 s rise time; 0.007 s fall time; 120 tones per sound; no gap between tones). Each stimulus sequence comprised 12 sets of tones and each set was 10 tones (0.4 seconds) long. The frequency of a tone could take on one of 70 values ranging from 700 to 2500 Hz (logarithmically spaced).

The selection of frequency values for each of the 12 sets depended on the stimulus condition. Four conditions were utilized: one condition contained no regularity (RAND), whereas the other three conditions (REG, REG2.5, and REG5) contained a regular pattern that started 2 seconds after sound onset. For the RAND condition (containing no regularity), 10 new frequency values were randomly selected for each of the 12 sets (Figure 1A, top). In the REG condition (containing a regularity), 10 new frequency values were randomly selected for each of the first 5 sets (0-2 seconds; similar to RAND), and then 10 new random frequency values were selected and repeated for set 6 to 12, thereby creating a regularity (the serial order of tone frequencies was identical in these repeating sets; Figure 1A, second row). Selection of the 10 random frequency values (and their serial order), used to create the regularity, differed from trial to trial. This condition is similar to the sounds used in previous studies that investigated sustained neural activity (Barascud et al., 2016; Southwell et al., 2017).

In order to investigate neural synchronization and sustained neural activity concomitantly, two additional conditions were utilized. Although the tone sequence repeats in the REG condition, the frequencies of the brief tones did not relate to each other in any regular way and the serial order of tone frequencies differed from trial to trial. Hence, the REG sequences may not drive observable oscillatory neural patterns, because observation of neural synchronization (e.g., using inter-trial phase coherence; ITPC) requires that neural activity is driven with the same oscillatory stimulus pattern on each trial (Henry and Obleser, 2012; Henry et al., 2014). Accordingly, we created regularities in which the tone sequences not only repeated (as in REG) but the change in frequency from tone to tone created a sinusoidal oscillatory pattern, to which synchronization of neural activity could be analyzed. We utilized two different oscillatory rates in order to assess the generalization of our stimuli. In the REG2.5 condition, the 10 frequency values (repeated for sets 6 to 12) were selected such that their arrangement reflected a 2.5-Hz sinusoidal frequency modulation (FM) (Figure 1A, third row). In the REG5 condition, the 10 frequency values (repeated for sets 6 to 12) were selected such that their arrangement reflected a 5-Hz sinusoidal FM (Figure 1A, bottom). For conditions REG2.5 and REG5, the starting phase was similar across trials, so that, when phase coherence is calculated, we would not be canceling signals with different phases. The precise frequency values chosen differed from trial to trial to avoid sounds being too similar (holding FM frequency and phase constant).

Throughout Experiment I, participants listened passively to 138 trials of each condition while watching a movie of their choice with subtitles and the sound muted on a battery-driven, portable DVD player. The experiment was divided into 6 blocks. During each block, 23 trials of each condition were randomly presented. Trials were separated by a 2-s inter-stimulus interval.

### EEG analysis

In order to investigate the sustained response, for each condition, the mean amplitude within the 2–4.8 second time window post stimulus onset and across the fronto-central-parietal electrode cluster was calculated. A one-way rmANOVA with the factor Condition (RAND, REG, REG2.5, REG5) was carried out using amplitude as a dependent measure.

Neural synchronization was investigated using ITPC calculated for neural activity within the 2–4.8 second time window. ITPC was averaged across 0.2-Hz wide frequency windows centered on the frequency in the stimulus (2.5 Hz; 5 Hz), and averaged across the fronto-central-parietal electrode cluster. Separately for ITPC values at 2.5 Hz and ITPC values at 5 Hz, a one-way rmANOVA with the factor Condition (RAND, REG, REG2.5, REG5) was carried out.

### Results

Figure 1B shows the response time courses and topographical distributions for each condition. The rmANOVA revealed a main effect of Condition (F_3,45_ = 5.52, p = 0.003, η_p_^2^ = 0.269). Post hoc tests indicated that responses were more negative in the 2–4.8 second time window for REG (t_15_ = −4.53, p_FDR_ = 0.01, r_e_ = 0.760), REG2.5 (t_15_ = −3.39, p_FDR_ < 0.05, r_e_ = 0.659), and REG5 (t_15_ = −2.93, p_FDR_ = 0.05, r_e_ = 0.603) compared to the RAND condition (Figure 1C). There were no statistically significant differences among the REG, REG2.5, and REG5 conditions (for all, p_FDR_ > 0.05). Scalp topographies indicate a wide distribution covering parietal electrodes as well as fronto-central electrodes.

Neural synchronization was statistically assessed for the ITPC at 2.5 Hz and at 5 Hz. The rmANOVA for ITPC values at 2.5 Hz revealed a main effect of Condition (F_3,45_ = 22.61, p = 9^e−6^, ε = 0.535, η_p_^2^ = 0.601). Post hoc tests showed significantly larger ITPC for REG2.5 (i.e., the regularity with a 2.5-Hz rhythm) compared to the RAND (t_15_ = 4.73, p_FDR_ = 0.001, r_e_ = 0.774), REG (t_15_ = 5.60, p_FDR_ < 0.001, r_e_ = 0.823), and REG5 conditions (t_15_ = 5.45, p_FDR_ < 0.001, re = 0.815).

The rmANOVA for ITPC values at 5 Hz revealed a main effect of Condition (F_3,45_ = 21.53, p = 4^e−6^, ε = 0.604, η_p_^2^ = 0.589). Post hoc tests showed significantly larger ITPC for REG5 compared to RAND (t_15_ = 4.06, p_FDR_ = 0.01, r_e_ = 0.724) and REG (t_15_ = 4.13, p_FDR_ < 0.01, r_e_ = 0.729), but also significantly larger ITPC for REG2.5 compared to RAND (t_15_ = 6.74, p_FDR_ = 0.001, r_e_ = 0.867) and REG (t_15_ = 7.25, p_FDR_ < 0.001, r_e_ = 0.882). The 5-Hz effect for the REG2.5 condition reflects an ITPC increase at the first harmonic frequency of 2.5 Hz. A large first harmonic in the EEG spectrum is a common observation for frequency-modulated sounds at low modulation rates (Picton et al., 2003). Scalp topographies are consistent with neural sources in auditory cortex (Näätänen and Picton, 1987; Picton et al., 2003).

Spearman correlations we calculated in order to investigate the relation between the sustained response effect for the REG condition (REG-minus-RAND) and the sustained response effect for the REG2.5 (REG2.5-minus-RAND) and the REG5 conditions (REG5-minus-RAND) (Figure 3). We observed a positive correlation between the REG regularity effect and the REG2.5 regularity effect (r = 0.524, p = 0.040), whereas the correlation between the REG regularity effect and the REG5 regularity effect was not significant (r = 0.238, p = 0.329), but showed a similar trend.

**Figure 3:**
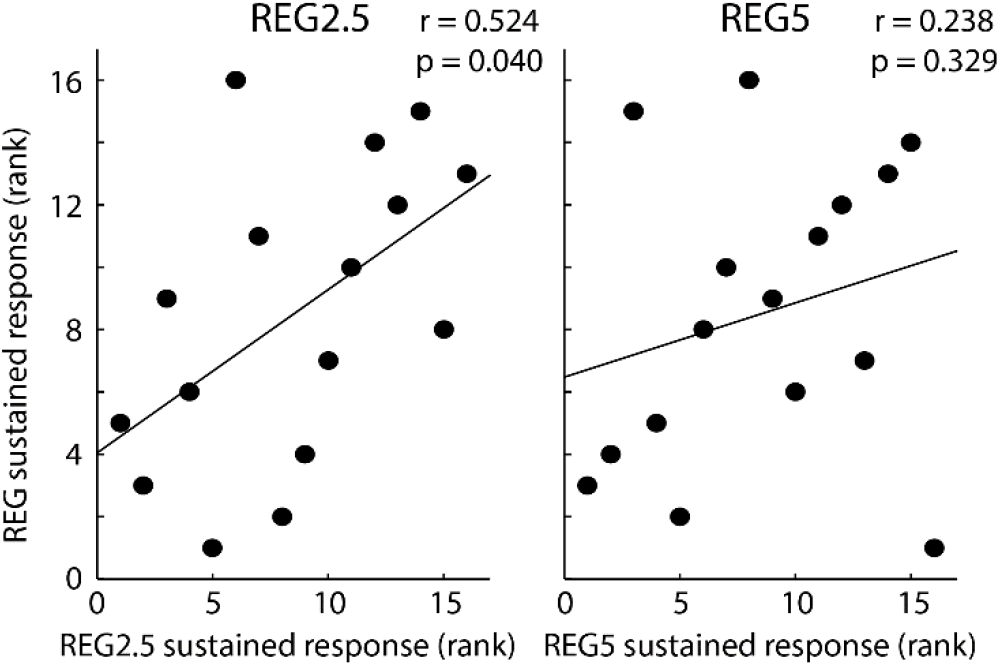
Spearman correlation between the sustained response effect for the REG condition versus the sustained response effect for the REG2.5 and REG5 conditions.

Repeated-measures correlation analyses were calculated (Bakdash and Marusich, 2017), in order to investigate the relation between the effect of regularity on sustained activity and ITPC for the REG2.5 and the REG5 conditions (Figure 4). To this end, we entered the two paired measures (RAND and REG2.5; RAND and REG5) of ITPC and sustained activity into the analysis. We observed negative correlations, indicating that the effect of regularity on sustained response increased (i.e., became more negative) when the effect of ITPC increased, which was significant for the REG2.5 condition (r = −0.603, p = 0.010), but not for the REG5 condition (r = −0.336, p = 0.187).

**Figure 4:**
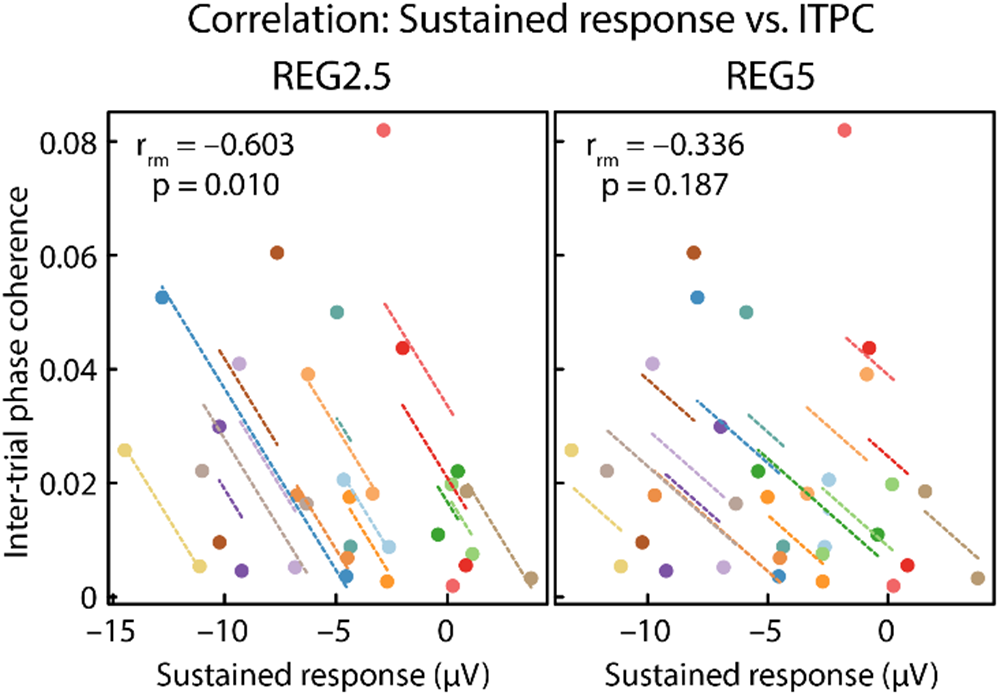
Repeated-measures correlations. Correlation between sustained response and inter-trial phase coherence. Note that the plots display negative correlations, because an increase in sustained activity reflects a negative-going signal, whereas an increase in ITPC is positive-going. Observations from the same participant are displayed in the same color, with the corresponding lines showing the fit of the repeated-measures correlation for each participant.

### Summary

The results of Experiment I replicate previous findings of a sustained response (here more negative) when a regularity occurs in a sound compared to when a sound contains no regularity (Barascud et al., 2016; Sohoglu and Chait, 2016; Southwell et al., 2017). In addition, Experiment I extends these previous reports by showing that, when the stimuli are constituted to allow observation of synchronization (REG2.5 & REG5), both the sustained response and neural synchronization are observed. The magnitudes of these signals correlate within participants, suggesting that they are not independent. In Experiment II we introduce a within-subject, parametric manipulation of regularity in order to observe whether both ITPC and the sustained response scale with the degree of regularity.

## Experiment II: Parametric modulation of neural synchronization and sustained activity

In order to parametrically modulate the strength of neural synchronization, we made use of narrowband sounds with varying degrees of frequency modulation (FM). Previous work shows that neural activity synchronizes with FMs in sounds (John et al., 2001; Herrmann et al., 2013; Henry et al., 2014) and manipulation of FM systematically modulates neural synchronization (Boettcher et al., 2002).

### Participants

Eighteen participants who did not participate in Experiment I participated in Experiment II (mean age [standard deviation]: 18.4 ±0.9 years; 10 female).

### Acoustic stimulation & procedure

Stimuli were narrow-band noises made by adding 48 frequency-modulated pure tones (components). For each of the 48 components, a random mean carrier frequency (Cf) between 1200 Hz and 2000 Hz was selected (starting phase was random). The range over which the carrier frequency was modulated was defined as Cf ±Cf×0.25 (e.g., for a mean carrier frequency of 1200 Hz, the carrier frequency was modulated between 900 Hz and 1500 Hz). For each of the 48 components, the rate at which the carrier frequency was modulated was not fixed but changed randomly between 2 Hz and 5 Hz within each component (the randomization differed between components). The 48 components were summed in order to obtain a narrow-band noise sound. We henceforth refer to this stimulus as the S0 condition (an abbreviation for a stimulus with no congruent frequency components; see Figure 5A, top).

**Figure 5:**
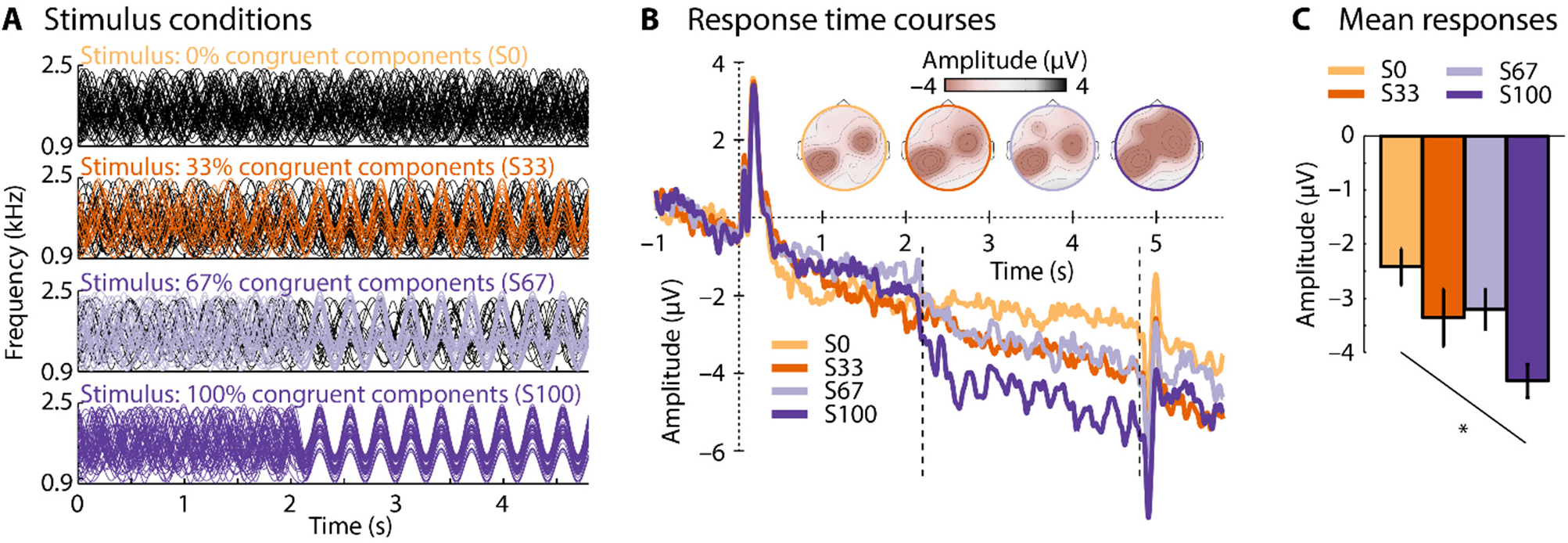
Stimulus conditions and neural responses for Experiment II. **A**: Examples of each of the sound conditions. Note that the representation of the auditory stimulus does not reflect the waveform of the narrow-band noise but instead the 48 frequency components (sound frequency on the y-axis). In S33, S67, and S100, 16, 32 or all 48 frequency components (respectively) become synchronized in phase at about 2.2 s after sound onset. **B**: Response time courses and scalp topographies for each condition. The dotted vertical lines mark the time window of interest used for analysis (2.2–4.8 s). **C**: Mean response magnitude for the 2.2–4.8 s time window. Condition labels: S0 – stimulus with 0% congruent frequency components; S33 – 33% congruent components; S67 – 67% congruent components; S100 – 100% congruent components. Error bars reflect the standard error of the mean (removal of between-subject variance; (Masson and Loftus, 2003). *p < 0.05.

Three additional conditions were created in which the degree of a sound’s FM phase coherence was manipulated. In the S33 condition, the FM rate of each of the 48 components changed randomly between 2 Hz and 5 Hz (similar to S0), but approximately 2.2 seconds after sound onset 16 out of the 48 frequency components (33%) synchronized in phase, and FM rate remained constant at 3.5 Hz for the rest of the sound (Figure 5A, second row). In the S67 condition, 32 out of the 48 frequency components (67%) synchronized in phase, and FM rate remained constant at 3.5 Hz for the rest of the sound (Figure 5A, third row). Finally, in the S100 condition, all 48 frequency components (100%) aligned in phase at about 2.2 seconds after sound onset and FM rate remained constant at 3.5 Hz for the rest of the sound (Figure 5A, bottom). Stimuli were created such that there was no transient change at the transition from nonaligned to phase-aligned frequency components.

Throughout Experiment II, participants listened passively to 138 trials of each condition while watching a movie of their choice with subtitles and the sound muted on a battery-driven, portable DVD player. The experiment was divided into 6 blocks. During each block, 23 trials of each condition were randomly presented. Trials were separated by a 2 s inter-stimulus interval.

### EEG analysis

The sustained response was analyzed by calculating the mean amplitude within the 2.2–4.8 second time window post stimulus onset and across the fronto-central-parietal electrode cluster. For each participant, a linear function fit was used to relate FM phase coherence (0, 33, 66, 100%) to the sustained response. In order test whether the sustained response was sensitive to FM phase coherence, the slope of the linear function was tested against zero using a one-sample t-test.

Inter-trial phase coherence (ITPC) was calculated for neural activity within the 2.2–4.8 second time window post stimulus onset. ITPC was averaged across a 0.2-Hz wide frequency window for two frequencies: one was centered on the stimulus’ modulation rate (3.5 Hz) and one on the first harmonic (7 Hz). We focused on these two frequencies as FM stimulation commonly leads to spectral peaks at the stimulation frequency and its first harmonic (Picton et al., 2003; Henry et al., 2017). ITPC was averaged across the fronto-central-parietal electrode cluster. Separately for each participant, and for ITPC values at 3.5 Hz and ITPC values at 7 Hz, a linear function fit was used to relate FM phase coherence (0, 33, 66, 100%) to ITPC. In order test whether ITPC was sensitive to FM phase coherence, the slope of the linear function was tested against zero using a one-sample t-test (separately for 3.5 Hz and 7 Hz).

### Results

Figure 5B shows the response time courses and topographical distributions for each condition. We observed a significant negative linear trend (t_17_ = −3.88, p = 0.001, r_e_ = 0.686; Figure 5C), indicating that the sustained response increased in magnitude (i.e., became more negative) as a function of FM phase coherence.

Figure 6 shows the ITPC frequency spectrum and topographical distributions for each condition. For the 3.5 Hz frequency, we observed a significant linear trend (t_17_ = 3.18, p = 0.005, r_e_ = 0.611), indicating that ITPC increased with increasing FM phase coherence. No effect of FM phase coherence was observed for the 7-Hz frequency (t_17_ = 1.10, p = 0.686, r_e_ = 0.258).

**Figure 6:**
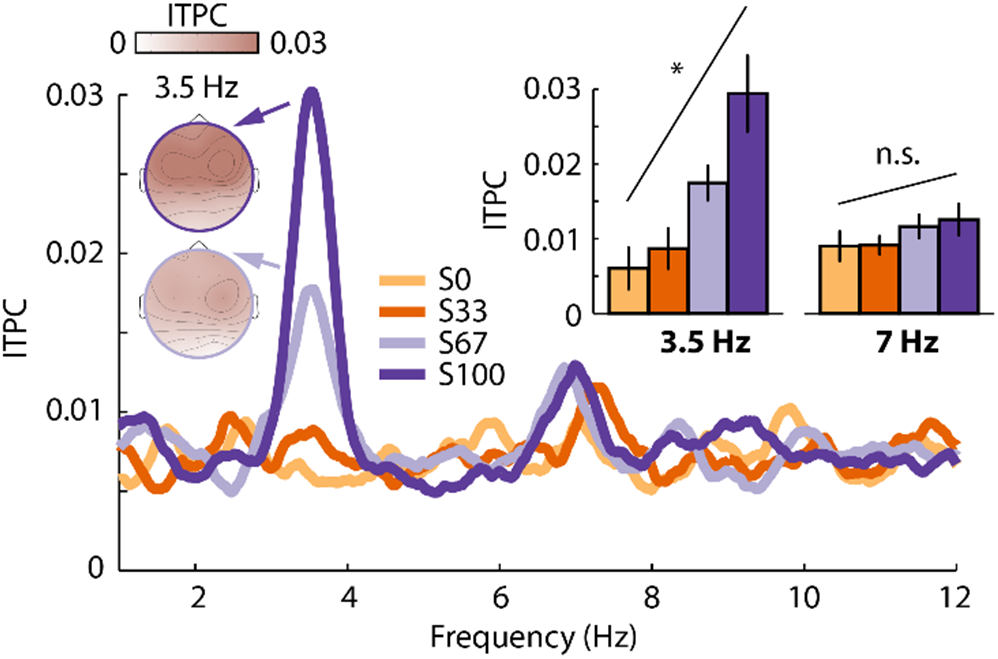
Inter-trial phase coherence (ITPC) results for Experiment II. Condition labels: S0 – stimulus with 0% congruent frequency components; S33 – 33% congruent components; S67 – 67% congruent components; S100 – 100% congruent components. Error bars reflect the standard error of the mean (removal of between-subject variance; (Masson and Loftus, 2003). *p < 0.05. n.s. – not significant

In order to compare the magnitude of the sustained response modulation with the magnitude of the ITPC modulation, the sustained response data and the ITPC data were separately z-transformed. The z-transformation was obtained as follows: We calculated the average response “AR” across all conditions and participants. The AR was subtracted from the response of each condition and participant. The resulting value for each condition and participant was subsequently divided by the AR. The sustained response data were also sign-inverted to ensure that for both ITPC and sustained activity, more positive values mean a larger response. Linear functions were fit to the z-transformed data as a function of FM phase coherence (0, 33, 67, 100%), and the slopes were compared between sustained response data and ITPC data using a paired t-test. This analysis revealed that neural synchronization (ITPC) was more strongly modulated by the degree of FM phase coherence in a sound compared to the sustained response (t_17_ = 2.45, p = 0.026, r_e_ = 0.511); in other words, the slope was steeper. The weaker apparent relation between the sustained, low frequency response and regularity may be owing to the sustained response being more sensitive to biological noise such as sweating, movement etc. compared to ITPC, which is an amplitude-normalized index. In order to account, in part, for such differences, we recalculated the analysis by applying a rationalized arcsine transform (Studebaker, 1985) to ITPC data before calculating the z-transformation. The rationalized arcsine transform linearizes indices that range from 0 to 1, such as ITPC, and makes them more Gaussian. Similar to the original analysis, ITPC was more strongly modulated by the FM phase coherence than was the sustained response (t17 = 3.62, p = 0.002, re = 0.660).

Finally, we investigated whether the systematic modulation of the sustained response by FM coherence is correlated with the modulation of ITPC using a repeated-measures correlation (Bakdash and Marusich, 2017). To this end, we entered the four paired measures (S0, S33, S67, S100) of ITPC and sustained activity into the analysis. We observed a negative correlation, indicating that an increase in sustained response (i.e., more negative) was accompanied by an increase in ITPC, but this correlation was only marginally significant (r = −0.263, p = 0.053; Figure 7).

**Figure 7:**
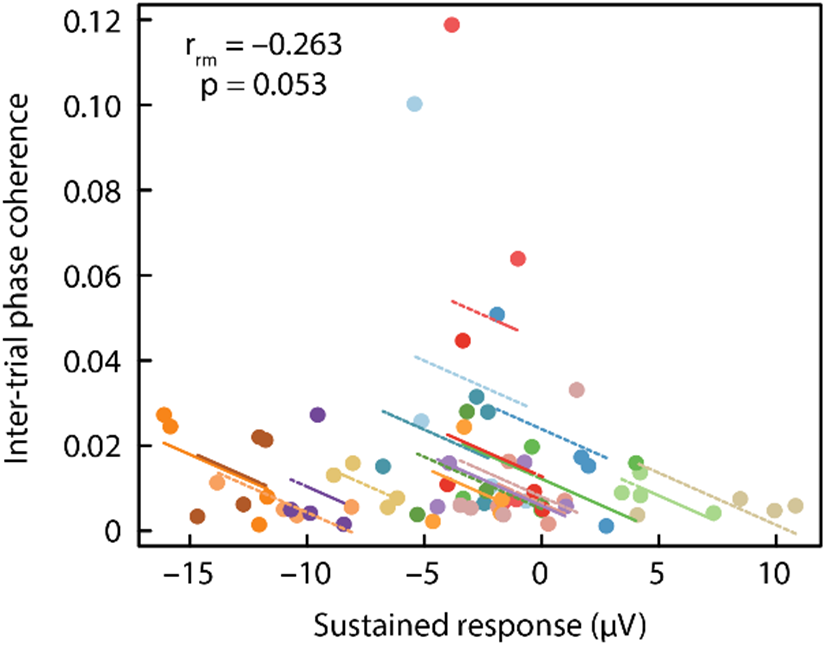
Repeated-measures correlation between sustained response and inter-trial phase coherence. Note that the plots display a negative correlation, because an increase in sustained activity reflects a negative-going signal, whereas an increase in ITPC is positive-going. Observations from the same participant are displayed in the same color, with the corresponding lines showing the fit of the repeated-measures correlation for each participant.

### Summary

The results of Experiment II show that parametrically manipulating the degree of frequency modulation in a sound leads to systematic changes in the magnitude of the sustained response and neural synchronization. The data also indicate that synchronization of neural activity is more sensitive to regularity imposed by a coherent frequency modulation in sounds compared to the sustained response. The data might thus indicate that the relation between neural synchronization and sustained activity levels is indirect, rather than that they are tightly linked. In order to investigate the relationship between the sustained response and neural synchronization further, we conducted Experiment III in which we utilized a selective attention task. If the sustained response and neural synchronization are differentially affected by attentional state, this would be evidence that the two neural systems are at least partially independent.

## Experiment III: Effects of attention on sustained response and neural synchronization

Experiment III aimed to investigate the degree to which attention affects the sustained response and neural synchronization. Participants either attended to sounds with and without regularity or ignored the sounds and attended to a visual multiple object tracking (MOT; (Pylyshyn and Storm, 1988) task that requires continuous attentional focus (Cavanagh and Alvarez, 2005; Alvarez and Franconeri, 2007; Tombu and Seiffert, 2008; Scholl, 2009) and is thus suitable for distraction over multiple seconds (Masutomi et al., 2016; Herrmann and Johnsrude, 2018).

### Participants

Twenty-seven participants who did not participate in Experiments I & II participated in Experiment III (mean age [standard deviation]: 20.3 ±3.3 years; 15 female). Three additional participants did not comply with the task instructions and were thus excluded.

### Acoustic stimulation & procedure

Acoustic stimuli were narrow-band noises similar to the stimuli used in Experiment II. In one condition, none of the 48 frequency components were synchronized in modulation rate over time. In a second condition, 38 out of 48 frequency components (79%) became synchronized at about 1.6 seconds after sound onset. The sound’s duration varied between 4 and 5 seconds (which was related to the duration categorization task described below).

During the experiment, participants sat in front of a Dell LCD computer screen (~70 cm away; 75 Hz repetition rate; 24-inch diagonal). On every trial, participants were concurrently presented with a visual moving dot display and an auditory stimulus (Figure 8A). Each trial started with a 1.3-s stationary display of 16 dots (dot diameter: 1.2 cm [0.9°]) of which 10 were white (distractor dots) and 6 were red (target dots). Presentation of dots was constrained to a display frame of 20.6 cm width (15.6°) and 19.4 cm height (14.7°) centered on the screen and highlighted to the participants by a gray frame on a black background. A yellow fixation square (0.16 cm [0.12°]) was presented at the center of the display frame. After 1.3 s, all dots reverted to white and the dots started to move. Concurrently with the dot movement and for the duration of the dot movement, one of the two sound conditions was played to the participant via headphones. Dots never moved outside of the display frame and never overlapped during movements; dots moved approximately 3.7 cm/s (2.8°/s). After 4 or 5 seconds (equal number of trials of each), the sound ended, the dot display froze, and one dot was marked in green (Figure 8A).

**Figure 8:**
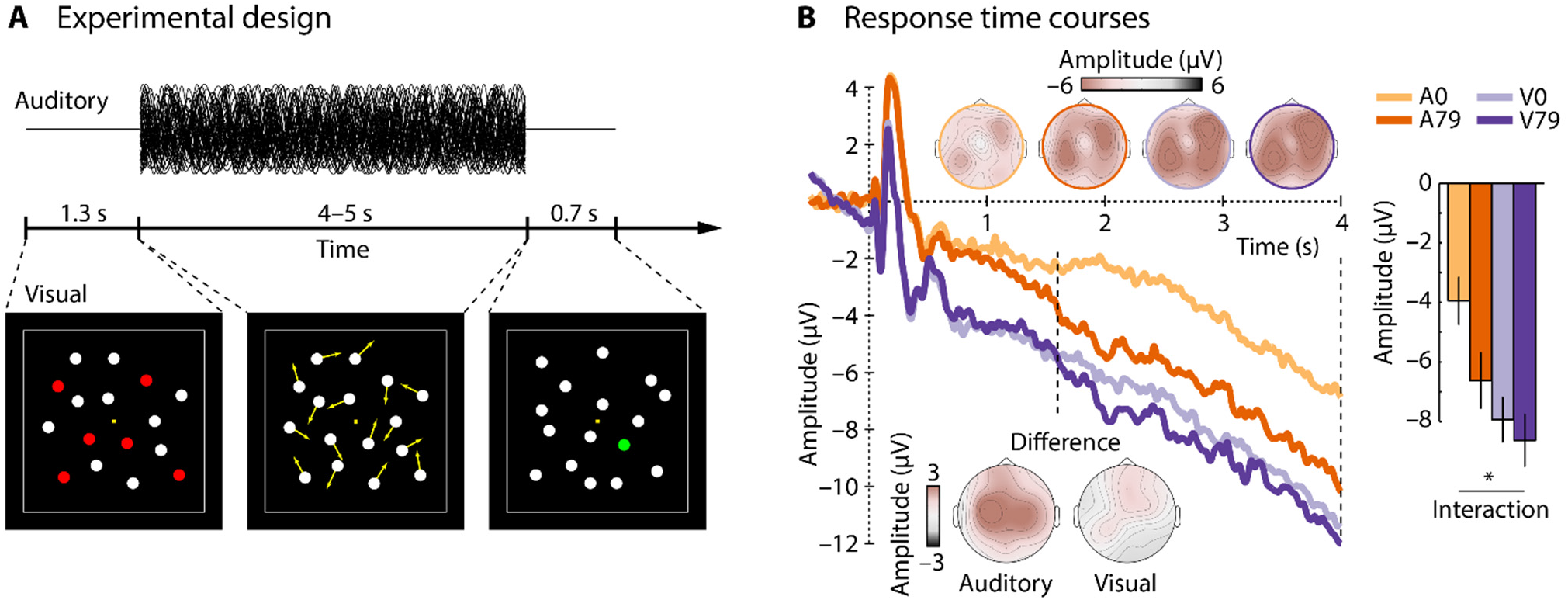
Experimental design and response time course for Experiment III. **A**: Shows the experimental design. Participants performed either an auditory duration categorization task (half of the blocks) or a visual multiple object tracking task (the other half of the blocks). Note that the representation of the auditory stimulus does not reflect the waveform of the narrow-band noise but instead the 48 frequency components similar to Figure 5A top. **B**: Response time courses and scalp topographies for each condition. The dotted vertical lines mark the time window of interest used for analysis (1.6–4 s). Bar graphs show the mean response magnitude for the 1.6–4 s time window. Condition labels: A0 – attention to auditory task, 0% congruent stimulus components; A79 – attention to auditory task, 79% congruent stimulus components; V0 – attention to visual task, 0% congruent stimulus components; V79 – attention to visual task, 79% congruent stimulus components. Error bars reflect the standard error of the mean (removal of between-subject variance; (Masson and Loftus, 2003). *p < 0.05.

Participants either performed a visual multiple object tracking (MOT) task or an auditory duration categorization task. In the MOT task, participants were asked to track those dots over time that were marked in red (target dots) in the pre-movement 1.3-s time period. They were also asked to ignore any sounds played concurrently with the dot movements. Participants had to judge whether the green dot was one of the dots they had been asked to track. In half of the trials, the green dot was indeed one of the target dots. In the other half of the trials, the green dot was one of the distractor dots. In the auditory duration categorization task, participants were asked to indicate whether they heard a sound with a short or long duration. During the auditory task, participants were asked to look at the fixation point on the screen, but to ignore the visual stimulation (which included the same MOT stimuli). Prior to the EEG recordings, participants underwent two training blocks, one for the visual task and one for the auditory task, and were taught what “short” and “long” durations were for the auditory task.

In order to target a d′ of about 2 (i.e., difficult but manageable) (Macmillan and Creelman, 2004) and to control for task difficulty differences between the visual task and the auditory task across participants, slight adjustments from the described procedure were implemented. The precise stimulation parameters were as follows: For 15 participants, the stimulus sound (and dot movements) lasted either 4 s (short) or 5 s (long) and, when the green dot was a distractor dot, it was selected *randomly* out of the 10 distractor dots. For 8 participants, the stimulus sound (and dot movements) lasted either 4 s (short) or 5 s (long) and, when the green dot was a distractor, it was chosen to be the distractor dot that was located *closest* to a target dot, making the MOT task more difficult. For 4 participants, the stimulus sound (and dot movements) lasted either 4.1 s (short) or 4.9 s (long), making duration categorization more difficult (as in variant 1 described above, green distractor dots were selected *randomly*). Critically, the auditory and visual stimulation was the same for all participants for the first four seconds after sound onset and we restricted our data analyses to this common period.

Participants performed 6 blocks of about 11–12 min each. During three blocks, participants performed the visual task; in the other three blocks they performed the auditory task. Task blocks alternated and starting block was counterbalanced across participants. Within each block, 36 sounds of each condition (0% congruent components and 79% congruent components) were pseudo-randomly presented. Overall, participants listened to 108 sounds of each condition, once while they attended to the sounds and performed the sound-duration categorization task and once while they attended to the visual stimulation and performed the MOT task. Henceforth we refer to sounds with 0% congruent components and sounds with 79% congruent components in the auditory attention task as A0 and A79, respectively. We refer to V0 and V79 for the same sounds presented during the visual attention task.

### Behavioral analysis

Behavioral data were analyzed using d′ (Macmillan and Creelman, 2004). For the visual task, when the participant responded “yes”, we counted the response as a hit if the green dot was indeed a target, and as a false alarm if it was not. For the auditory task, when the participant responded “long”, we counted the response as a hit if the sound duration was indeed long (5 s or 4.9 s) and as a false alarm if it was not.

A few participants struggled with one or both of the tasks, exhibiting relatively poor performance. In order to ensure that we included only those participants who, we were confident, were attending to the instructed task most of the time, we excluded participants with poor task performance from subsequent analyses. Hence, participants with a d′ < 1 (corresponding approximately to 0.7 proportion correct responses and lower; with chance level being 0.5), in either of the tasks, were excluded at this stage (N=6), leading to twenty-one participants that were included in the EEG analysis.

### EEG analysis

The sustained response was analyzed by calculating the mean amplitude within the 1.6–4 second time window and across the fronto-central-parietal electrode cluster. A two-way rmANOVA with the factors FM coherence (0% vs. 79% congruent frequency components) and Attention (auditory, visual) was carried out. In order to account for potential differences in task difficulty on the individual level between the auditory task and the visual task, the difference between the auditory d′ and the visual d′ was entered as a covariate of no interest into the analysis.

Inter-trial phase coherence (ITPC) was calculated for the responses within the 1.6–4 second time window. ITPC was averaged across 0.2-Hz wide frequency windows centered on the stimulus’ modulation rate and the first harmonic (3.5 Hz; 7 Hz), and averaged across the fronto-central-parietal electrode cluster. Separately for ITPC values at 3.5 Hz and ITPC values at 7 Hz, a two-way rmANOVA with the factors FM coherence (0% vs. 79% congruent frequency components) and Attention (auditory, visual) was carried out, again entering the difference between the auditory d′ and the visual d′ as a covariate of no interest.

### Results

Behavioral performance measured as d′ did not differ significantly between the auditory duration categorization task and the visual MOT task (t20 = 1.28, p = 0.215, r_e_ = 0.275; auditory d′ [±std]: 2.13 ±0.66; visual d′ [±std]: 1.90 ±0.60).

EEG response time courses and scalp topographies are displayed in Figure 8B. The rmANOVA revealed a main effect of FM coherence (F_1,19_ = 7.13, p = 0.015, η_p_^2^ = 0.273) and a marginally significant effect of Attention (F_1,19_ = 3.05, p = 0.097, η_p_^2^ = 0.138). Critically, the interaction between FM coherence × Attention was significant (F_1,19_ = 4.63, p = 0.044, η_p_^2^ = 0.196), due to a larger response difference between stimulus types when participants attended to auditory stimuli (sound duration task) compared to when participants attended to visual stimuli (MOT task). In fact, the sustained response was larger (more negative) for stimuli with 79% congruent frequency components compared to stimuli with 0% congruent frequency components only when participants attended to the auditory stimuli (F_1,19_ = 9.52, p_FDR_ = 0.05, η_p_^2^ = 0.334), but not when they performed the visual distraction task (F_1,19_ = 0.74, p_FDR_ > 0.05, η_p_^2^ = 0.037). An exploratory investigation of the effect of regularity on EEG activity separately for each electrode revealed only one electrode that was sensitive to auditory regularity when listeners performed the visual task (i.e., the sounds were not the focus of attention; Fp2; p = 0.027, uncorrected). In contrast, 11 out of the 16 electrodes were sensitive to regularity when listeners performed the auditory task (i.e., attended to the sounds; all p ≤ 0.05, uncorrected; Fp1, Fp2, Fz, F3, C3, C4, Cz, P3, P4, Pz, Oz).

Results for the inter-trial phase coherence (ITPC) analysis are shown in Figure 9. The rmANOVA for the 3.5-Hz frequency revealed a main effect of FM coherence (F_1,19_ = 12.40, p = 0.002, η_p_^2^ = 0.395), due to larger ITPC values for stimuli with 79% congruent frequency components compared to stimuli with 0% congruent frequency components. Neither the main effect of Attention (F_1,19_ < 0.01, p = 0.970, η_p_^2^ < 0.001) nor the FM coherence × Attention interaction were significant (F_1,19_ = 1.66, p = 0.213, η_p_^2^ = 0.080).

**Figure 9:**
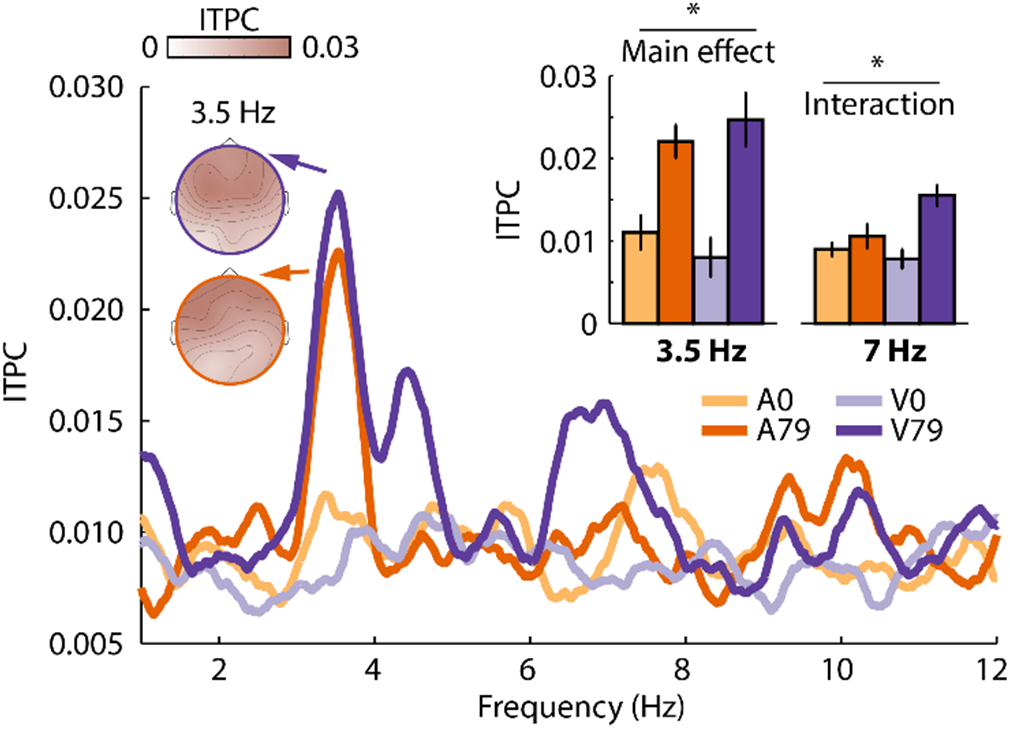
Inter-trial phase coherence (ITPC) results for Experiment III. Condition labels: A0 – attention to auditory task, 0% congruent stimulus components; A79 – attention to auditory task, 79% congruent stimulus components; V0 – attention to visual task, 0% congruent stimulus components; V79 – attention to visual task, 79% congruent stimulus components. Error bars reflect the standard error of the mean following removal of between-subject variance (Masson and Loftus, 2003). *p < 0.05.

The rmANOVA for the 7-Hz frequency (first harmonic) revealed a main effect of FM coherence (F_1,19_ = 9.82, p = 0.005, η_p_^2^ = 0.341), no effect of Attention (F_1,19_ = 0.81, p = 0.381, η_p_^2^ = 0.041), and a significant FM coherence × Attention interaction (F_1,19_ = 7.21, p = 0.015, η_p_^2^ = 0.275). The interaction was due to a larger ITPC difference between stimulus types when participants attended to the visual stimuli (MOT task) compared to when participants attended to the auditory stimuli (sound duration task). In fact, ITPC was larger for stimuli with 79% congruent frequency components compared to stimuli with 0% congruent frequency components only when participants attended to the visual stimuli (F 1,19 = 16.06, p_FDR_ = 0.05, η_p_^2^ = 0.458), but not when they attended to auditory stimuli (F_1,19_ = 1.46, p_FDR_ > 0.05, η_p_^2^ = 0.071).

### Summary

The results of Experiment III show a differential effect of attention on the regularity-related sustained activity compared to neural synchronization. The sustained activity was affected by attention, such that an effect of FM phase coherence (i.e., temporal regularity) was only observed when participants attended to the sounds. In contrast, neural synchronization at the stimulation frequency (3.5 Hz) was not affected by the manipulation of attention, whereas neural synchronization at the first harmonic (7 Hz) differed between stimulus conditions only when participants attended to the visual stimulation. These results show that, for the processing of temporal regularity in sounds, neural synchronization and sustained activity are differently affected by a distracting visual task.

Rather unexpectedly, we observed a marginally significant increase in sustained activity levels when participants performed the visual compared to the auditory task (Figure 8B). Note that, in conducting this study, we matched the behavioral performance level of the two tasks across participants, and accounted for any remaining within-subject differences between tasks by using the behavioral performance difference as a co-variate in our neural response analysis. This reduces the likelihood that the overall increase in sustained activity reflects a difference in task difficulty. It is also unlikely that the sustained response reached a ceiling level, given that it continued to increase (i.e., became more negative) throughout the epoch. Finally, another possibility is that visual activity increased when participants performed the MOT task (even though visual and auditory stimulation was physically identical in both tasks), leading to an overall increase in sustained activity. Although there have been reports of sustained activity for visual stimuli (Järvilehto et al., 1978), topographical distributions of the sustained response were very similar across the three experiments (without clear contributions from occipital electrodes), which suggests no additional generator in visual areas specifically in Experiment III compared to Experiments I and II.

## Discussion

The current set of experiments investigated the relation between two types of neural responses – synchronization and sustained activity – that are associated with the processing of temporal regularity in sounds. Experiment I demonstrates that neural activity synchronizes with a sound’s temporal regularity while sustained activity increases. In Experiment II, parametric manipulation of FM phase coherence in sounds systematically modulated neural synchronization and sustained neural activity, albeit neural synchronization was more sensitive to coherent frequency modulation. Experiment III revealed that under visual distraction neural activity synchronized with a sound’s temporal regularity, whereas sustained activity was not sensitive to this regularity. The current data show that, in the context of temporal regularity in sounds, neural synchronization and sustained neural activity co-occur, but that the two neural signatures are differentially affected when a listener is distracted by a demanding visual task.

### Sensitivity of neural synchronization and sustained activity to temporal regularity in sounds

In each of the three EEG experiments, we observed that neural activity synchronized with the temporal regularity (i.e., frequency modulation) in the sounds (Figures 2, 6, and 9). Synchronization of neural activity – sometimes referred to as phase-locking or entrainment of neural oscillations – is commonly observed for sounds that contain temporal regularity (John et al., 2002; Lakatos et al., 2008; Stefanics et al., 2010; Ding and Simon, 2012; Lakatos et al., 2013b; Kayser et al., 2015; Presacco et al., 2016; Herrmann et al., 2017), including frequency-modulated sounds similar to the ones utilized here (Boettcher et al., 2002; Picton et al., 2003; Herrmann et al., 2013; Henry et al., 2017).

The current data also show that, for sounds that consist of brief tones and narrow-band noise sounds, sustained neural activity increases following the onset of a regularity; in this case, coherent frequency modulation (Figures 5 and 8). Hence, our data are consistent with previous studies that have demonstrated sensitivity of sustained neural activity to other types of regularities in sounds such as repeating tone-sequence patterns and coherent frequency changes in broadband chords (Barascud et al., 2016; Sohoglu and Chait, 2016; Teki et al., 2016; Southwell et al., 2017). Our results are also in line with studies that report sustained activity in response to sounds with a very simple regular structure such as pure tones (Köhler and Wegner, 1955; David et al., 1969; Picton et al., 1978; Weise et al., in press).

Parametric manipulation of the degree of coherent frequency modulation (i.e., temporal regularity) in a narrow-band noise led to a systematic increase in both neural synchronization (Figure 6) and sustained neural activity (Figure 5) although neural synchronization was more sensitive to the degree of FM coherence compared to sustained neural activity. Within-participant correlations further indicate that the increase in neural synchronization and the increase in sustained activity are somehow related. Future work will investigate whether the neural populations underlying neural synchronization and sustained activity are directly coupled or whether their apparent relation is instead due to a third process that influences both.

One important contribution of the current study is that we demonstrate the simultaneous occurrence of increased sustained neural activity and neural synchronization for temporally regular sounds. Previous studies that investigated neural synchronization (e.g., (Maiste and Picton, 1989; Henry and Obleser, 2012; Herrmann et al., 2013; Keitel et al., 2017) have not reported changes in sustained activity, likely because the sustained activity is a low-frequency (DC – direct current) signal and the application of a high-pass filter is a common pre-processing step for electrophysiological data. The current data thus indicate that multiple, concurrent neural signals index the processing of temporally regular structure in sounds.

### The influence of attention on neural synchronization and sustained activity

In Experiment III, we showed that neural activity thought to originate in auditory cortex (Näätänen and Picton, 1987; Picton et al., 2003; Herrmann et al., 2013) synchronizes with a sound’s temporal regularity independent of attention. In contrast, the regularity-based increase in sustained response was only observed when participants attended to auditory stimuli or when they were lightly distracted by watching a movie (Experiments I and II), but not when participants were distracted by a demanding visual task (Experiment III).

Previous work investigating the neural correlates of attention to a sound’s rhythm (Elhilali et al., 2009; Xiang et al., 2010; Lakatos et al., 2013a; Zion Golumbic et al., 2013) observed that synchronization depended on attention. However, attention to a sound’s rhythm may enhance the representation of periodicity (and thus synchronization), whereas attention to a sound’s duration (as in this study) may not have the same effect. Consistent with our and previous work, much of the existing research suggests that detection of regularities in sounds can occur automatically (Sussman et al., 2007; Winkler et al., 2009; Bendixen, 2014; Sohoglu and Chait, 2016; Southwell et al., 2017; Sussman, 2017), but that attention may interact with automatic processing depending on the task (Sussman, 2017).

Previous work suggested that an increase in sustained activity due to a sound’s regularity is independent of attention (Sohoglu and Chait, 2016). In this study, an increase was observed when participants performed either a visual image-detection task (while ignoring sounds) or an acoustic change-detection task (i.e., attending to sounds). The difference between this previous work and our results may be due to two reasons. First, in the previous study, attention was manipulated using a between-subject design, in which one participant group performed the visual task and another group the auditory task (Sohoglu and Chait, 2016). In the current study, the same participants performed both the visual and the auditory task (in different blocks using a within-subject design), which may increase sensitivity. Second, in the previous study, images for the visual distraction task were presented at intervals ranging from 0.5 to 4 s (and only 20% of the images required a response), which may have left time for participant to switch attention to the auditory stimuli (Sohoglu and Chait, 2016). The MOT task utilized here requires continuous attention (Cavanagh and Alvarez, 2005; Scholl, 2009) and is thus suitable for distraction from auditory stimuli over multiple seconds (Masutomi et al., 2016). The data indicate that a demanding visual task may be suppressing regularity-based sustained activity rather than that attention to sounds is enhancing it.

### Neural synchronization and sustained activity may reflect different stages of regularity processing

The scalp topographies in the current study show a focal spatial distribution for neural synchronization as opposed to the more wide-spread spatial distribution for the sustained response. Source localization studies suggest the main source underlying neural synchronization to low-frequency periodicities (<8 Hz) is located in auditory cortex (Herrmann et al., 2013; Herrmann et al., 2016; Keitel et al., 2017), whereas the sustained activity appears to additionally involve the inferior frontal cortex, parietal cortex, and hippocampus (Barascud et al., 2016; Teki et al., 2016). A difference in neural sources between the two responses may also indicate a functional difference. Neural synchronization is directly related to stimulus acoustics as it reflects a neural signal that tracks a sound’s periodicity (temporal regularity). In contrast, an increase in sustained activity appears to be independent of a specific regularity in a sound as it has been observed for repeating tone-sequence patterns, coherent frequency changes in broadband chords, and here for frequency-modulated sounds (Barascud et al., 2016; Sohoglu and Chait, 2016; Teki et al., 2016). Hence, it is possible that neural synchronization may reflect a hierarchically lower, sensory process, whereas sustained neural activity reflects a more abstract process related to more general structure in sounds.

The results from Experiment III are consistent with the neural representation of a sound’s temporal regularity in auditory cortex being unaffected by attention, whereas an attentionally demanding visual task suppresses the neural representation of the regularity in higher-level brain regions. This is also in line with the well-accepted view of auditory processing that assumes that neural activity in hierarchically lower regions in the auditory system is mostly sensitive to acoustic properties of sounds and less receptive to a listener’s attentional state, whereas neural activity in hierarchically higher regions is more sensitive to the attentional state of a listener (whether a listener attends to auditory stimuli or ignores them; (Davis et al., 2007; Wild et al., 2012; Puvvada and Simon, 2017; Holmes et al., in press). We speculate that a listener’s attentional state affects the progression of the neural representation of a sound’s temporal regularity from auditory cortex to higher-level brain regions, and that this progression is suppressed in situations with distracting visual stimulation.

## Conclusions

The current study investigated the functional relationship between two neural signatures that index the detection of a regularity in sounds: neural synchronization and sustained activity. In three EEG experiments we show that neural synchronization and sustained activity occur concomitantly, that both neural signatures are parametrically modulated by the degree of temporal regularity in sounds, but that the two signatures are differentially affected when a listener is distracted by a demanding visual task. Our data may indicate that neural synchronization reflects a more sensory-driven response to regularity compared with sustained activity, which may be influenced by attentional, contextual, or other experiential factors.

## Acknowledgements

Research was supported by the Canadian Institutes of Health Research (MOP133450 to I.S. Johnsrude), and the CFREF BrainsCAN postdoctoral fellowship awarded to B. Herrmann. We thank Youngkyung Jung and Suvarna Moharir for their help during data acquisition.

